# Balancing the risks of mating: biogeographic evidence of cleistogamy as a bet hedging strategy

**DOI:** 10.1101/2024.03.28.587200

**Authors:** Maya Weissman, Dafeng Zhang, Rebecca Kartzinel, Daniel Weinreich

## Abstract

Cleistogamy is a mating system in which plants produce some proportion of closed, autonomously self-pollinating flowers. Cleistogamous flowers differ from chasmogamous flowers, which are open flowers capable of outcrossing. Both dimorphic cleistogamy (cleistogamous and chasmogamous flowers produced on the same plant) and complete cleistogamy occur. Cleistogamy has been hypothesized to be a bet hedging strategy for reducing risk in the face of unpredictable pollinator availability. However, conflicting results across species and challenges connecting theory to data have prevented researchers from proving that cleistogamy is bet hedging. To test the bet hedging hypothesis, we investigated the distribution of over 400 cleistogamous species through biogeographical analyses. We find that cleistogamy is more prevalent in cooler, more variable environments. Additionally, we find that among cleistogamous species, complete cleistogamy is more likely to occur in warmer, more stable, tropical and subtropical environments. We hypothesize that the difference in distribution between complete and dimorphic cleistogamy may be driven by the opposing forces of selection to increase cleistogamy proportion and extinction risk, which we test using a heuristic Markov transition model. We conclude that the distribution of cleistogamy suggests that the strategy has evolved in variable environments, consistent with expectations for bet hedging.

## 1 Introduction

Cleistogamy is a mating system in angiosperms where plants produce some proportion of closed, autonomously self pollinating flowers (Darwin [1862], Darwin [1897], Schemske [1978], Campbell et al. [1983], Berg and Redbo-torstensson [1998], Culley and Klooster [2007]). Cleistogamous flowers (literally ”closed marriage”) are in opposition to chasmogamous (”open marriage”) flowers, which are open, showy flowers capable of outcrossing (Darwin [1862], Darwin [1897], Schemske [1978], Campbell et al. [1983], Berg and Redbo-torstensson [1998], Culley and Klooster [2007]). Cleistogamy is believed to have evolved to reduce risk in the face of adverse environments, specifically in relation to variability in pollinator abundance (Schoen and LLoyd (1984), Berg and Redbo-torstensson [1998], Culley and Klooster [2007], Oakley et al. [2007]). Cleistogamy has been observed in 228 genera and 50 families of angiosperm, and is estimated to have evolved over 40 times (Culley and Klooster [2007]). The majority of cleistogamous species engage in dimorphic cleistogamy, or the production of both cleistogamous and chasmogamous flower morphs on the same plant. However complete cleistogamy, or the exclusive production of cleistogamous flowers, has also been observed (Culley and Klooster [2007]).

Cleistogamy has been hypothesized to be a bet hedging strategy (Schoen and LLoyd (1984), Berg and Redbo-torstensson [1998], Culley and Klooster [2007]). Bet hedging is a broad class of adaptations where a population reduces risk in the face of unpredictable environments (Slatkin [1974], Philippi and Seger [1989], Simons [2011]). Mathematically, bet hedging is defined as a trade-off where a population lowers its mean fitness in order to also reduce fitness variance across environments. While bet hedging has been observed across the tree of life, from mammals to bacteria, one of the best studied taxa within bet hedging research is the angiosperm clade (Simons [2011]). Other examples of bet hedging traits in angiosperms include delayed seed germination (Venable (1985), Venable (2007), Childs et al. [2010]), variance in germination phenology (Arthur et al. [1973]), production of multiple seed types through amphicarpy (Zhang et al. [2020]), and investment in a few, high fitness fruit via a high ovule:fruit ratio (Aker (1982)). In order to prove that a trait is bet hedging, one must demonstrate that the trait yields the expected trade-off in fitness mean and variance, and thus that the trait is adaptive in the environmental context in which it has evolved. Despite the fact that other bet hedging strategies in annual plants are well understood (especially delayed seed germination (Childs et al. [2010]), there has been limited research to confirm whether cleistogamy actually meets criteria for adaptive bet hedging.

Bet hedging strategies fall into one of two categories: conservative or diversified (Simons [2011]). Both conservative and diversified bet hedging strategies are observed amongst cleistogamous species. Conservative bet hedging is defined by the production of a single generalist, low-risk phenotype. While the conservative phenotype is expected to have a lower fitness than a specialist phenotype when environmental conditions are good, the conservative phenotype will have a higher fitness in stress conditions (Slatkin [1974], Simons [2011], Starrfelt and Kokko [2012]). We thus consider complete cleistogamy to be form of conservative bet hedging. Cleistogamous flowers can be considered a low-risk, conservative phenotype because they do not require biotic pollinators, and therefore will experience no variability in fertilization success due to pollinator availability. Additionally, the closed flower morphology provides other fitness benefits: they are energetically inexpensive to produce, and often have higher fitness due to increased fruit/seed set (Schemske [1978], Schoen and LLoyd (1984), Oakley et al. [2007]). On the other hand, diversified bet hedging is a strategy where an isogenic lineage produces multiple different phenotypes. The advice ”don’t put all your eggs in one basket” is often used to visualize the utility of diversified bet hedging. This ensures that some proportion of offspring will always be well adapted, at the cost of sacrificing some proportion of maladapted offspring (Slatkin [1974], Simons [2011], Starrfelt and Kokko [2012]). Dimorphic cleistogamy is thus a diversified bet hedging strategy, defined by the production of two flower phenotypes. Dimorphic cleistogamous species get the benefits of producing some proportion of cleistogamous flowers, while also gaining the benefits of occasional outcrossing through the production of chasmogamous flowers. The main benefit of chasmogamy is thought to be the production of genetically variable offspring via outcrossing, which reduces inbreeding and extinction risk, as well as potentially leading to heterosis (Berg and Redbo-torstensson [1998], Culley and Klooster [2007]). However, the chasmogamous flowers on dimorphic cleistogamous plants can, and frequently do, also self (Oakley et al. [2007]). In summary, the differences in morphology between cleistogamous and chasmogamous flowers have two consequences, both of which can be considered to reduce risk: probability of selfing and energy economy. In this way, dimorphic cleistogamy is different from the more common method of mixed mating in angiosperms, which utilize monomorphic flowers capable of both selfing and outcrossing (Goodwillie et al. [2005]). Dimorphic cleistogamy has been hypothesized to be more adaptive than monomorphic mixed mating when selection to maintain outcrossing is stronger (Oakley et al. [2007]).

Theoretical models of cleistogamy have invoked environmental heterogeneity and bet hedging to explain selection for cleistogamy (Berg and Redbo-torstensson [1998], Culley and Klooster [2007], Schoen and LLoyd (1984), LLoyd (1984), Waller [1984]). These models assume that each flower type is adapted to a different environmental condition, and fluctuations in environment allow for the maintenance of a mixed strategy (Schoen and LLoyd (1984), LLoyd (1984)). Previous research has established that cleistogamy tends to be associated with stressful environments (Culley and Klooster [2007], Oakley et al. [2007], Campbell et al. [1983]). However, determining which environmental conditions are optimal for each flower morph has been challenging and often contradictory. For example, increased cleistogamous flower production has been associated with decreases in temperature (Lord [1982], Oakley et al. [2007]), but also increases in temperature (Austin et al. [2022]); long days (Langer and Wilson [1965]), but also short days (Heslop-Harrison [1962]); higher density (Cheplick and Quinn [1983]), but also no relationship with density (Stølen and Shands [1975]); increased plant size (Joseph Trapp and Hendrix [1988]), but also decreased plant size (Berg and Redbo-torstensson [1998]). Further, there have been opposing observations in which seasons cleistogamous and chasmogamous flowers are favored. Many species produce the two flower morphs sequentially across different seasons, rather than simultaneously. Yet again, no consistent pattern has emerged: some species produce the cleistogamous flowers first (Oakley et al. [2007], De Clavijo (1997), Cheplick and Clay [1990]), others produce the chasmogamous flowers first (REDBO-TORSTENSSON and Berg [1995], Jasieniuk and Lechowicz [1987], Bell and Quinn [1987]), and still others produce both flowers simultaneously (Le Corff [1993], Schoen [1984]). As such, more work is needed to determine large-scale trends in the types of environmental stressors that select for cleistogamy, and how this translates to the maintenance of complete and dimorphic cleistogamy.

Another obstacle in explaining the maintenance of dimorphic cleistogamy has been estimating the relative fitness of the two flower morphs. In the majority of studies, the fitness advantage of cleistogamous flowers is so high that the maintenance of chasmogamy seems disadvantageous, even when accounting for inbreeding depression (Oakley et al. [2007], Winn and Moriuchi [2009], Albert et al. [2011]). Oakley et al. [2007] found that cleistogamous flowers can have up to a 231% fitness advantage compared to chasmogamous flowers. Although again, there are exceptions to this rule. In some species, both cleistogamous and chasmogamous flowers have equal fitness (Culley [2002]), and in others chasmogamous flowers have the higher fitness (Koontz et al. (2017)).

Finally, while the primary benefit of cleistogamy is thought to be assured fertilization in the face of variability in biotic pollinator (e.g. insect) abundance, cleistogamy has been observed in over 300 species of wind pollinated grasses(Campbell et al. [1983], Culley and Klooster [2007]). In fact, cleistogamy is more common in the Poaceae family than any other angiosperm family, with approximately 5% of grass species exhibiting the strategy(Campbell et al. [1983]). Despite the fact that variability in pollinator abundance is irrelevant to abiotic pollinated species, studies of cleistogamy in grasses have still linked its evolution to environmental variability, with the majority of cleistogamous species described as colonizers of disturbed habitats or drought stress tolerators (Campbell et al. [1983]). Thus, it seems that cleistogamous flowers may provide benefits in the face of environmental stressors other than pollinator availability.

Given the large diversity in cleistogamous species and challenges connecting theory to data, an outstanding question in the field has been identifying large-scale trends in cleistogamy evolution and evaluating whether cleistogamy can be considered a bet hedging strategy. Here, we utilize biogeographical analyses to investigate the distribution of cleistogamy in order to evaluate the bet hedging hypothesis. We find that overall, cleistogamy is more prevalent in cooler, more variable environments, consistent with expectations for bet hedging. Additionally, we contrast differences amongst species that engage in cleistogamy. We first contrast members of Poaceae, or wind pollinated grasses, with all other species as a proxy for pollination mode. We find that non-Poacea species are more prevalent in colder environments, where biotic pollinator abundance is the most variable (Ollerton [2017], Baskett et al. [2020]). We also find significant differences in the distribution of complete vs. dimorphic cleistogamy, regardless of pollinator mode. Amongst occurrences of cleistogamy, dimorphic cleistogamy is associated with cooler, more variable environments, while complete cleistogamy is more likely to occur in warmer, more stable, tropical and subtropical environments. We hypothesize that the difference in distribution between complete and dimorphic cleistogamous strategies may be driven by the opposing forces of selection to self and extinction risk, which we test using a heuristic Markov transition model. Our results suggest that, despite the large diversity in cleistogamous species, large-scale patterns in the distribution of cleistogamy reveal that the strategy has evolved in variable environments, consistent with bet hedging predictions.

## 2 Materials and methods

### 2.1 Data and code availability

Data and code are available at: https://github.com/mweissman97/cleistogamy_biogeography/.

### 2.2 Occurrence data for cleistogamous species

We began by taking a list of 628 cleistogamous species and their respective strategy (i.e. complete vs. dimorphic) from Culley and Klooster [2007].

We then generated our dataset of occurrences. First, we found all geographic occurrences of each species by querying the Global Biodiversity Information Facility (GBIF) (GBIF.org (2024)). Occurrences were filtered so that only those with exact coordinates and a basis of record of preserved specimen (i.e. museum or herbaria collections) were used to control for quality. Each coordinate was classified into one of four latitudinal zones based on the absolute value of their latitude: tropic (*<* 23.5^◦^), subtropic (23.5^◦^ − 35^◦^), temperate (35^◦^ − 66.5^◦^), and polar (*>* 66.5^◦^) (Britannica [2024], Britannica [a], Britannica [b], Society [2012]). Coordinates were also used to determine the ecosystem of the plant using the World Wildlife Federation (WWF) terrestrial ecoregions of the world (Olson et al. [2001]). Coordinates were classified into one of 827 ecoregions, which are then grouped into one of fourteen bioclimatic zones, or biomes. Before filtering, there were 8,0951 occurrences across 628 species.

Next, we filtered our occurrence data using the Kew Royal Botanical Gardens Plants of the World Online (Govaerts et al. (2021), Brown et al. [2023]). First, species names were reconciled. Second, geographic occurrences were checked against the known native ranges of the species. Points outside of the native range or within the invasive range of the species were removed, in order to focus on only the environmental context where species evolved. We note that there is significant sampling bias in herbaria records, which is reflected in our dataset (Daru et al. [2018]). Therefore, we summarize species occurrence within each ecoregion by taking the mean of all species occurrence coordinates within said ecoregion. By only counting each species occurrence within an ecoregion once, we correct for oversampling due to relative abundance and collection bias. Post filtering steps, we were left with 5,277 occurrences of 425 unique species.

Finally, we connected coordinates to bioclimatic data using WorldClim (Hijmans et al. [2005]). Climate data was downloaded at a resolution of 2.5 arc minutes, which corresponds to approximately 4.5 km at the equator.

#### 2.2.1 **Statistical analyses**

For differences in presence vs. absence (both across latitudinal zones and biomes), we utilized the test of equal or given proportions. The test of equal proportions asks whether the proportion of occurrences is significantly different between groups, taking into account differences in the size of each group (Newcombe [1998], Team (2010)). For differences in climate between groups, we utilized a two sample t-test to analyze differences in mean. P-values were adjusted using the Benjamini-Hochberg procedure to control for false positives and correct for multiple testing (Benjamini and Hochberg [1995], Team (2010)).

Data querying and filtering, as well as subsequent data visualization and analyses were carried out using the R programming language (Team (2010)).

### 2.3 Markov transition model

We develop a simple Markov transition model to explore potential drivers of the difference in latitudinal distribution of complete vs. dimorphic strategies. In our model, can adopt one of three discrete strategies: complete chasmogamy, dimorphic cleistogomay, or complete cleistogamy. We assume species start by adopting the completely chasmogamy strategy, as this has been identified as the ancestral state (Culley and Klooster [2007]). Species can then evolve to increase their cleistogamous flower production. Transitions in only this direction are possible due to the immediate fitness advantage of cleistogamy; seeds from cleistogamous flowers are almost always larger and more fit than those from chasmogamous flowers (Oakley et al. [2007]). Species transition from complete chasmogamy to dimorphic cleistogamy, and then from dimorphic cleistogamy to complete cleistogamy. Therefore, species cannot transition from complete chasmogamy to complete cleistogamy in a single time step. Finally, due to inbreeding, complete cleistogamous species go extinct.

We mathematically express this model in a Markov transition matrix, which specifies the probability of transitioning from any starting state to any other state in one time step. Species transition from the complete chasmogamy state (*CH*) to the dimorphic state (*Dim*) at rate *c*, or the cleistogamy transition rate. Species in the dimorphic cleistogamy state can also increase their cleistogamous flower production at the same rate *c*, thereby entering the complete cleistogamy state (*CL*). Finally, complete cleistogamous species go extinct at rate *e*. When a species exits the complete cleistogamy state due to extinction, it is replaced by a new complete chasmogamous species, in order to keep the total number of species constant (see fig. 4A). The transition matrix is thus:

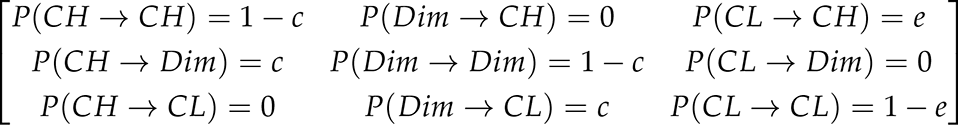

The expected frequency of each state at equilibrium was estimated by computationally calculating the stationary distribution of the transition matrix. The stationary distribution (*π*) is the matrix that satisfies the condition *π* = *π* × *T*.

Matrices were constructed across the range of all possible values of *c* and *e* at a step size of 0.01. Matrix multiplication was carried out using the MatLab Programming Language (MATLAB (2010)).

#### 2.3.1 **Latitudinal zone simulations**

We developed a model to test the hypothesis that the differences in frequency of both complete and dimorphic strategies between latitudinal zones may be due to the opposing pressures of selection to increase cleistogamy proportion and extinction rate. We assume that moving away from the equator (i.e. from tropical to temperate) increases both selection to increase cleistogamy, as pollinators decrease and environments become more variable (Geppert et al. [2023], Ludewig et al. (2023), Baskett et al. [2020], Ollerton [2017]), and also extinction rate (Mittelbach et al. [2007], Saupe [2023], Weir and Schluter [2007]). We wanted to investigate which values of *c* and *e*, given the above constraints, yield the observed patterns in the latitudinal gradient of the distribution of both complete and dimorphic cleistogamy. Namely, that complete cleistogamy is most prevalent in the subtropics, and that dimorphic cleistogamy increases in frequency moving away from the equator. As such, we performed replicate simulations where random points in the *c* and *e* parameter space that matched all criteria were selected from each latitudinal zone. Two thousand replicate simulations were carried out.

Because one of our constraints is that we assume both *c* and *e* must increase as the absolute value of latitude increases, the space of possible parameters for each latitudinal zone depends on the parameters chosen for the previous zone. Therefore, we begin by selecting the parameter value pair for the tropical zone. First, all possible points in parameter space that could correspond to the tropic zone were determined based on the observed frequency of complete and dimorphic cleistogamy from our earlier biogeographic analysis (*Dim <* 0.45 and *CL*/(*Dim* + *CL*) *<* 0.25). Simulations began by selecting one of these points at random and assigning it as the tropic parameters. We next assign the parameter values for the subtropic zone, based on the constraints given by the tropic parameters. The space of possible subtropic parameter values is constrained by the value of the tropic parameter values (*c_subtropic_ > c_tropic_*and *e_subtropic_ > e_tropic_*). Additionally, given that the realized frequency of both complete and dimorphic cleistogamy increases, we also constrain possible subtropic parameters to those in which the absolute frequency of both complete and dimorphic cleistogamy increases relative to the tropic parameters. The parameter values for the subtropic zone is then selected randomly from all possible points. Simulations failed to assign parameters to the subtropic zone when the *c* and *e* parameter values for the tropic point were so high that no possible points in parameter space could meet all criteria. In those cases, the parameter values randomly assigned for the tropic zone were outside of the possible range for our simulations, and those point were skipped, which occurred in roughly half of simulations. This allowed us to experimentally find the upper limits of the range of possible *c* and *e* values for the tropic zone. Finally, the point in parameter space corresponding to the temperate zone is assigned. Similar to before, we assume that both *c* and *e* must increase relative to the subtropic parameter values. Relative to the subtropic point, we also constrain possible parameters to those where the absolute frequency of dimorphic increases and the absolute frequency of complete decreases. The parameter values for the temperate zone is then chosen randomly from all possible points. In this phase of the simulations, we never failed to find parameter values that met these criteria.

Simulations and visualization were carried out using the R programming language (Team (2010)).

## 3 Results

### 3.1 Cleistogamy is more common in variable environments

We utilize the terrestrial ecoregions of the world (Olson et al. [2001]) to explore the places where cleistogamy is present and absent. Ecoregions are distinct assemblages of natural communities, such as ”Northeastern US coastal forests.” We determined if any cleistogamous species occurrences were within the borders of each of the 827 ecoregions in order to visualize occurrences on a map (fig. 1). Ecoregions were labelled ”absent” if no cleistogamous species occurred within the ecoregion. Ecoregions were labelled ”present” if one or more cleistogamous species occurred within the ecoregion. For ecoregions where cleistogamy is present, in the map we further subdivided the ecoregions based on which strategies occurred: ”complete only”, ”dimorphic only”, and ”both” (fig. 1).

**Figure 1:**
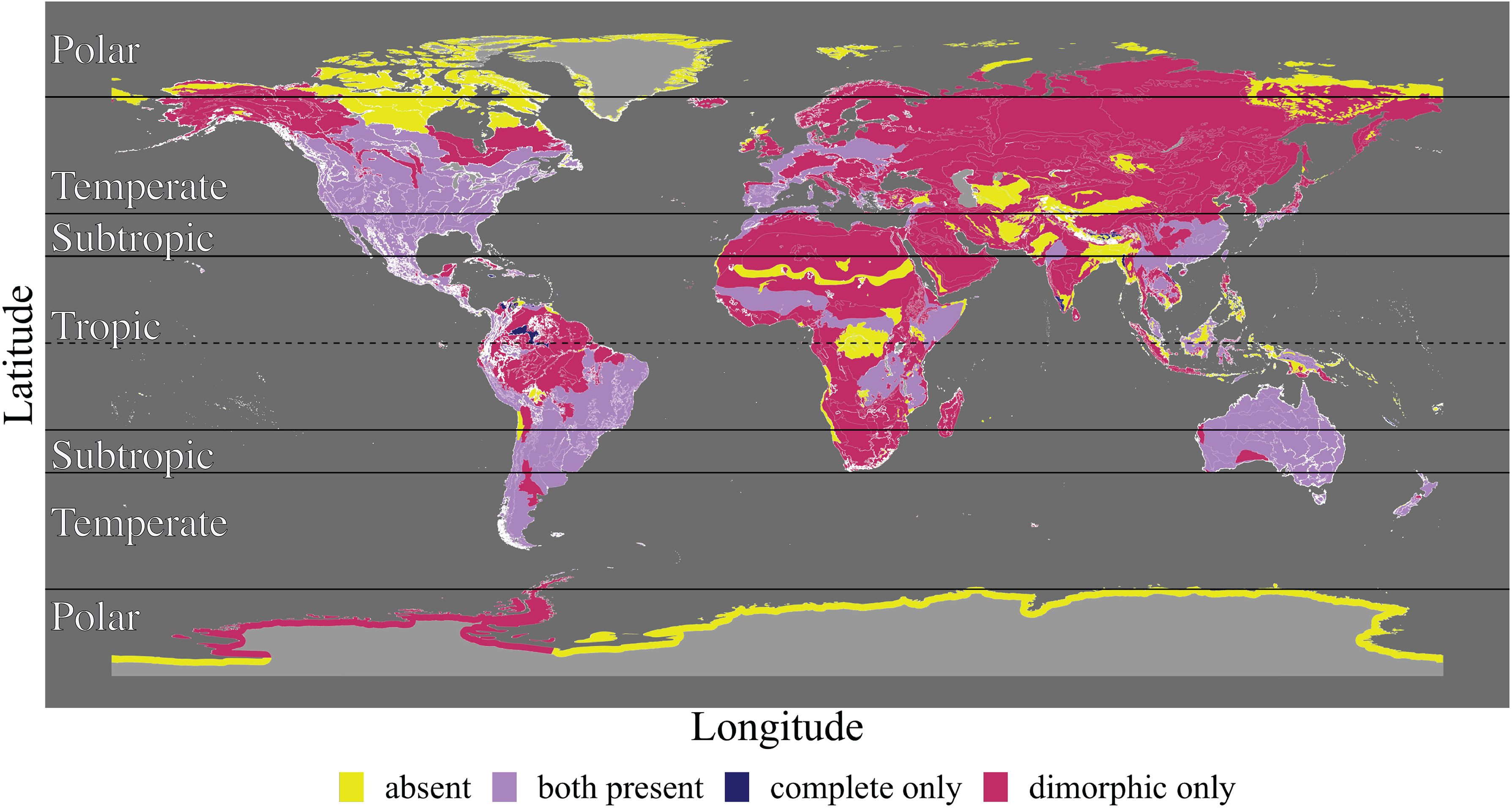
The earth is subdivided into 827 distinct terrestrial ecoregions (Olson et al. [2001]), colored by whether cleistogamy is absent (yellow) or present. Ecoregions where cleistogamy is present are further classified by the type of cleistogamous strategy present: both complete and dimorphic strategies (purple), only complete cleistogamous species (blue), and only dimorphic cleistogamy species (pink). The dotted line shows the equator. Polygons for which no ecoregion data exists (e.g. lakes, ice caps) are filled in with dark grey. Ecoregion polygon borders are shown in white. Horizontal lines denoting the latitudinal zones are also shown, from equator to poles: tropic, subtropic, temperate, and polar.

In order to further analyze our occurrence data, we first focus on contrasting ecoregions in which cleistogamy is present and absent, irrespective of cleistogamous strategy, to investigate the ecoregions and climates where cleistogamy occurs. Cleistogamy is thought to reduce risk in the face of unpredictable biotic pollinator abundance, which has been shown to be correlated with temperature (Ollerton [2017], Baskett et al. [2020]). Therefore, if cleistogamy is bet hedging, we would expect cleistogamy presence to be associated with cooler, more unpredictable environments.

We find that cleistogamy is significantly more prevalent in the subtropic and temperate latitudes than the other latitudinal zones (*p <* 10^−9^), which are associated with cooler mean temperatures and larger climate variability (Society [2012]) (fig. 2A). Conversely, cleistogamy is more likely to be absent in tropic (or constantly warm) and polar (constantly cold) latitudes (Britannica [a], Britannica [b], Britannica [2024]). Ecoregions were sorted into one of four latitudinal zones (tropical, subtropical, temperate, and polar) based on the latitude of the centroid of the ecoregion. The overall proportion of ecoregions where cleistogamy is present (74%) is denoted with the black line.

**Figure 2:**
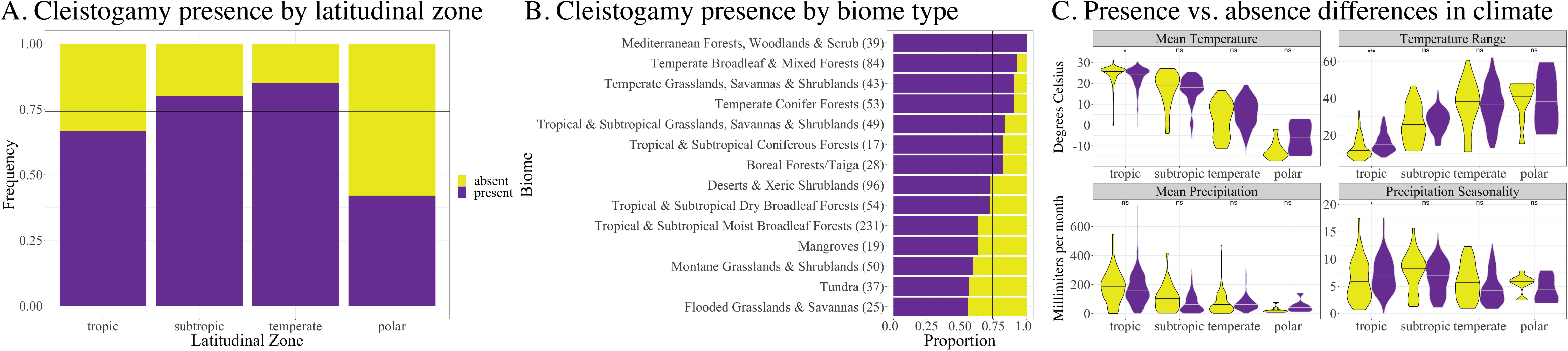
Ecoregions were classified by whether cleistogamy was absent (yellow) or present (purple). A. Ecoregions were subdivided by the absolute value of the latitude of the centroid into 4 zones: tropic (*<* 23.5), subtropic (between 23.5 and 35), temperate (between 35 and 66.5), and polar (*>* 66.5). The overall proportion of ecoregions where cleistogamy is present is denoted with a black horizontal line. Cleistogamy is more prevalent in subtropical and temperate latitudes, and absent more in tropic and polar latitudes. B. Ecoregions were classified by their biome type, with the number of ecoregions within each biome presented in parentheses. Again, the overall proportion of ecoregions where cleistogamy is present is denoted with a black vertical line. C. Differences in climate between ecoregions where cleistogamy is present vs. absent by latitudinal zone. Only within the tropic zone are there significant differences in climate between ecoregions where cleistogamy is present vs. absent. Within the tropic zone, cleistogamy is present in ecoregions with a lower mean temperature, wider temperature range, and higher precipitation seasonality. Differences in mean precipitation were not significant.

We can also observe this pattern by investigating the relative frequency of cleistogamy presence across biomes (fig. 2B). Again, the overall proportion of ecoregions where cleistogamy is present is denoted with the black line (74%). Biomes are not independent of latitudinal zone (i.e. temperate broadleaf forests are found exclusively in temperate latitudes), however some biomes are found in multiple latitudinal zones. Compared to all other biomes, cleistogamy is significantly more prevalent in Mediterranean forests, woodlands and scrub (*p* = 0.004) and temperate broadleaf and mixed forests (*p* = 0.001). Both of these biomes can be considered variable environments, with large seasonal shifts in climate in addition to high between-year variance in environment (Rundel et al. [2016], Dreiss and Volin [2020], Loidi et al. (2022)). For example, Mediterranean ecoregions are highly diverse ”mosaic climates” defined by their hot and dry summers and cold and wet winters. Mediterranean ecoregions also have high between-year climate variance, as the biome has both high drought and wildfire risk (Rundel et al. [2016]). Temperate broadleaf forests also exhibit distinct warm and cool seasons, with moderate annual average temperatures. Fire disturbances are also common in temperate broadleaf forests (Dreiss and Volin [2020]). Conversely, cleistogamy is absent significantly more in tropical and subtropical moist broadleaf forests compared to all other biomes (*p <* 10^−4^). Tropical and subtropical moist broadleaf forests are a ”constantly good” biome, characterized by low variability in annual temperature and high levels of rainfall (Atangana et al. (2014), Loidi et al. (2022)).

Finally, we investigated differences in climate between ecoregions where cleistogamy is present or absent, controlling for latitudinal zone (fig. 2C). We focus on four climatic variables: mean temperature (degrees Celsius), temperature range (degrees Celsius), mean precipitation (millimeters per month), and precipitation seasonality (millimeters per month). These climatic variables were selected to summarize both mean and spread for temperature and precipitation. Mean temperature, mean precipitation, and precipitation seasonality are all also relatively uncorrelated with each other (*corr <* 0.5), while temperature range is only correlated with mean temperature (*corr* = −0.73) (supplemental fig. 1A). However, temperature range is especially interesting to us, because within-year temperature range is correlated with between-year temperature variability (supplemental fig. 1B).

Within the tropic zone, cleistogamy is present in ecoregions with lower mean temperature (*p* = 0.033), wider temperature range (*p* = 0.0002), and higher precipitation seasonality (*p* = 0.032). (fig. 2C). Climate differences were only significant within the tropic zone. We saw no significant difference in mean precipitation within the tropic zone (*p* = 0.32). All p-values presented are adjusted to correct for multiple testing. For a full table of all nineteen bioclimatic variables and their respective p-value in each latitudinal zone, see Supplemental Table 1.

### 3.2 Environment correlates with differences in the distribution of type of cleistogamy strategy

In order to investigate differences in distribution amongst cleistogamous species, we now compare species occurrence, rather than cleistogamy presence and absence. For every ecoregion, we determine all species present. We summarize multiple species occurrences by taking the mean latitude and longitude coordinates of all species, rather than the centroid of the ecoregion. This allows for slightly more fine grained climate analyses. In this framing, the same ecoregion can be counted more than once in our dataset, as more than one species may be found in the same ecoregion. Conversely, the same species can also appear multiple times, as one species can be found in multiple ecoregions.

We contrast the spatial distributions of the two types of cleistogamy strategies: complete and dimorphic. We also contrast species that are members of the Poaceae family with all other species. Members of the Poaeceae family, or grasses, are wind pollinated (Campbell et al. [1983]). The remaining taxa are primarily biotic (such as insect) pollinated. We thus use phylogeny as a proxy for pollinator mode, to determine where cleistogamy may be reducing risk of variability in pollinator availability, and where cleistogamy may be correlated with other environmental stressors.

It has previously been observed that dimorphic cleistogamy is far more prevalent than complete cleistogamy, with 87.8% of cleistogamous species adopting the dimorphic strategy (Culley and Klooster [2007]). We find that there is a great deal of geographic overlap in the distribution of complete and dimorphic cleistogamy; both strategies are present in over 40% of ecoregions where cleistogamy has been observed at least once (fig. 1). However, we do see some differences in the biogeography of the two cleistogamy strategies. First, complete cleistogamy is most prevalent in the subtropical zone, while dimorphic cleistogamy is most prevalent in the temperate latitudinal zone (fig. 3A). Another way to visualize the latitudinal gradient is to investigate the frequency of both strategies, rather than absolute counts (fig. 3B). The overall proportion of dimorphic cleistogamous species is denoted with the black line (0.88). We now see that the proportion of species that adopt the complete strategy is not significantly different between the tropic and subtropic zones (*p* = 0.792). The proportion of complete species is significantly lower in the temperate zone than in the tropical and subtropical zones (*p <* 10^−16^).

**Figure 3:**
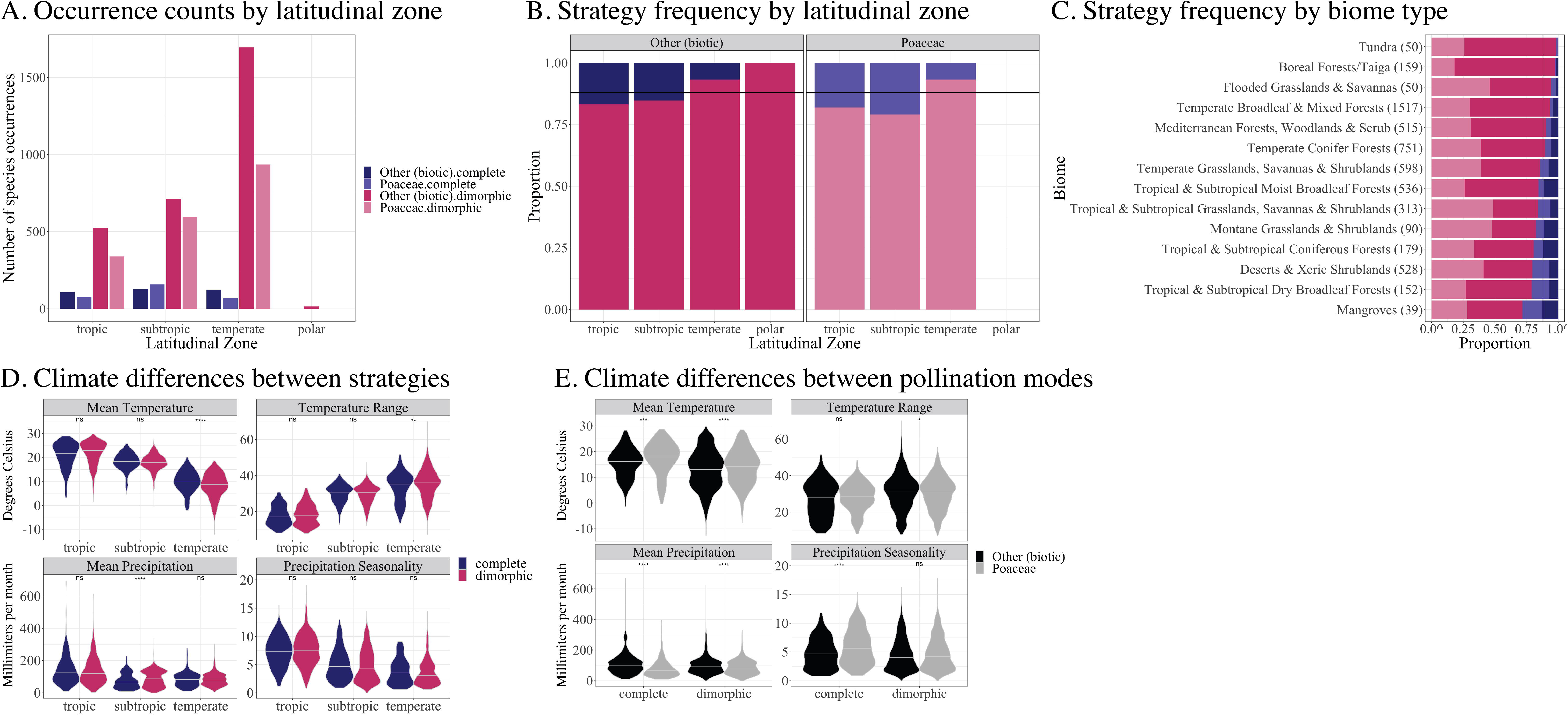
A. Count of species occurrences per ecoregion by latitudinal zone. We subdivide by both cleistogamy strategy (complete vs. dimorphic) and pollination mode (Poaceae Family vs. Other Families). The number of occurrences of the dimorphic strategy increases moving from tropic to temperate zones. The number of occurrences of the complete strategy is at its maximum in the subtropic zone. The prevalence of Poaceae species is lowest in the temperate zone. B. Strategy frequency by latitudinal zone, subdivided by pollination mode. The overall proportion of dimorphic species is denoted with the black line (0.88). Complete cleistogamy is more prevalent in tropic and subtropic zones, while dimorphic cleistogamy is more prevalent in the temperate zone. Complete cleistogamy is more prevalent among Poaceae (0.138) than among other species (0.109). C. Occurrences by biome type, with the number of occurrences within each biome presented in parentheses The overall proportion of occurrences of dimorphic cleistogamy is shown with the black vertical line. D. Differences in climate between occurrences of complete vs. dimorphic by latitudinal zone, irrespective of pollination mode. Complete cleistogamy is found in climates with significantly higher mean temperate and more narrow temperature ranges within the temperate zone. Within the subtropic zone, complete cleistogamy occurs in climates with significantly lower mean precipitation. E. Differences in climate between Poaceae and members of other families by cleistogamy strategy, irrespective of latitudinal zone. Other, primarily biotic pollinated, species occur in climates that have a lower mean temperature and higher mean precipitation. Among complete cleistogamous species, non-Poaceae species are associated with lower precipitation seasonality.

We also investigate differences between members of Poaceae, or wind pollinated grasses, and all other families, or primarily biotically pollinated species. Overall, complete cleistogamy is significantly more prevalent amongst Poaceae than all other families (*p* = 0.0011) (fig. 3A). Members of Poaceae are also less prevalent than other families in the temperate zone (*p <* 10^−9^) (fig. 3A). However, the relative distribution of complete and dimorphic cleistogamy across latitudinal zones within each pollinator mode are the same (*p* = 1) (fig. 3B).

We next investigate differences in distribution by biome (fig. 3C). We tested whether the proportion of species that adopts each strategy significantly differs between biomes. The overall proportion of species with dimorphic cleistogamy (88%) is again shown by the black line. Dimorphic cleistogamy is significantly more prevalent in boreal forests (*p* = 0.003) and temperate broadleaf and mixed forests (*p <* 10^−12^) compared to all other biomes. These are two of the coldest biomes, with the largest seasonal variations in climate (Loidi et al. (2022), Saucier et al. [2015], Dreiss and Volin [2020]). As colder weather is correlated with lower biotic pollinator abundance, it follows that cleistogamy will be favored in these biomes. Additionally, high seasonality in climate will likely favor the sequential, instead of simultaneous, production of different flower morphs, which is common in species that engage in dimorphic cleistogamy. Complete cleistogamy is significantly more prevalent in mangroves (*p* = 0.037), tropical and subtropical dry broadleaf forests (*p* = 0.009), deserts and xeric shrublands (*p <* 10^−8^), and tropical and subtropical coniferous forests (*p* = 0.025) compared to all other biomes. These are all warmer biomes on average, which is consistent with our previous finding that complete cleistogamy is more prevalent in tropical and subtropical latitudes (Loidi et al. (2022)). Both mangroves and deserts have been described as variable environments, with high variance in both temperature and precipitation (Quinn (2008), Huntley (2023)). This aligns with the theory that complete cleistogamy may be a form of conservative bet hedging. Still, it is surprising that complete cleistogamy is significantly less prevalent in biomes where increased cleistogamous flower production is expected to be favored. This suggests that there are additional disadvantages to complete cleistogamy, especially in colder, more variable biomes.

We also find differences in biome distribution by pollinator mode. We find that non-members of Poaceae, or primarily bitoic pollinated species, are significantly more prevalent in boreal forests / taiga (*p <* 10^−6^), temperate broadleaf and mixed forests (*p <* 10^−9^), and tropical and subtropical moist broadleaf forests (*p <* 10^−4^). In colder biomes (i.e. boreal forests, temperate forests, and tundra), it is unsurprising that cleistogamy will be more prevalent in biotic pollinated species, as colder mean temperatures are correlated with decreased biotic pollinator abundance (Ollerton [2017], Baskett et al. [2020]). However, it is interesting that cleistogamy is more prevalent in non-Poaceae species in tropical and subtropical moist broadleaf forests, given this biome is characterized by a consistently warm and wet climate (Loidi et al. (2022), Atangana et al. (2014)). This may be due to differences in the distribution of grasses irrespective of cleistogamy, as the thick canopy of tropical rainforests limits the successes of grasses (Atangana et al. (2014)). In contrast, members of Poaceae are significantly more prevalent in deserts and xeric shrublands (*p <* 10^−11^), which tend to be more variable in precipitation than temperature (Quinn (2008)). Unsurprisingly, members of the grass family are also more prevalent in all types of grassland biomes than other biomes: montane (*p* = 0.042), temperate (*p* = 0.015), and tropical and subtropical (*p <* 10^−9^).

We next investigated differences in climate between the complete and dimorphic strategies. We control for latitudinal zone, but temporarily ignore pollination mode (fig. 3D). We find that complete cleistogamy is more prevalent in milder, less variable environments than dimorphic cleistogamy. Within the temperate zone, complete cleistogamy tends to occur in places with warmer mean temperatures (*p <* 10^−4^) and more narrow temperature ranges (*p* = 0.009). We also find that complete cleistogamy is associated with lower mean precipitation within the subtropic zone (*p <* 10^−5^). We find no significant differences in precipitation seasonality between the two strategies. For the p-values of differences in mean for all nineteen bioclimatic variables, see Supplemental Table 2.

We now focus on differences in climate between members of the Poaceae family and other species, controlling for cleistogamy strategy (fig. 3E). Given the similar latitudinal gradients (fig. 3B), we ignore latitude zone. Non-Poaceae species are associated with lower mean temperature (*p <* 10^−4^). Among only dimorphic cleistogamous species, non-Poaceae species are also associated with wider temperature ranges (*p* = 0.0467). Since biotic pollinator relative abundance is correlated with mean temperature (Ollerton [2017], Baskett et al. [2020]), it follows that pressure to evolve cleistogamy will increase in biotic pollinated species in ecoregions with colder mean temperature and higher temperature variance. Members of the Poaceae family are associated with lower mean precipitation (*p <* 10^−5^). Among only complete cleistogamous species, species in Poaceae are associated with higher precipitation seasonality (*p <* 10^−5^). This suggests that cleistogamy may reduce risk against precipitation stresses in wind pollinated species. For the p-values of all nineteen bioclimatic variables, see Supplemental Table 3.

### 3.3 Modeling the latitudinal gradient in complete vs. dimorphic cleistogamy strategy

Regardless of pollinator type, we see a strong pattern in the distribution of complete and dimorphic cleistogamy: complete cleistogamy is favored in milder, more stable environments (i.e. subtropic latitudes), while dimorphic cleistogamy is more prevalent in cooler, more variable environments (i.e. temperate latitudes). Why does the overall number of cleistogamous species increase, while the proportion of complete cleistogamy decreases, as environments become colder and less stable? One possible hypothesis is that colder, more variable environments pose an additional challenge to complete cleistogamy due to an increased risk of extinction. Moving away from the equator leads environments to become colder and less reliable, thereby increasing the selective pressure to engage in cleistogamy driven by decreased pollinator availability (Geppert et al. [2023], Ludewig et al. (2023), Baskett et al. [2020], Ollerton [2017]). However, moving toward the poles will also lead extinction risk to increase (Mittelbach et al. [2007], Saupe [2023], Weir and Schluter [2007]). As complete cleistogamy leads to high levels of inbreeding, this makes those species especially susceptible to extinction risk.

In order to test this hypothesis, we developed a simplified Markov transition model to predict the distribution of strategies (fig. 4A). We simplify and consider only three strategies, or states: complete chasmogamy (*CH*), dimorphic cleistogamy (*Dim*), and complete cleistogamy (*CL*). We assume that transitions in only one direction (to increase cleistogamy rate) can occur, given the immediate fitness advantage of cleistogamy. Therefore, species transition from complete chasmogamy to dimorphic at a rate *c*, or the selective pressure to increase cleistogamy proportion. Species transition from dimorphic to complete cleistogamy at the same rate *c*, as we assume that the selective advantage of increasing cleistogamy proportion is constant. In other words, we assume that increasing cleistogamy proportion from 0.0 to 0.1 offers the same fitness advantage as increasing cleistogamy proportion from 0.9 to 1.0. Finally, complete cleistogamous species go extinct at rate *e*, and are replaced with new chasmogamous species, keeping the total number of species constant. Using this framework, the Markov model can then numerically estimates the proportion of species in each state as a function of the transition rates between states.

**Figure 4:**
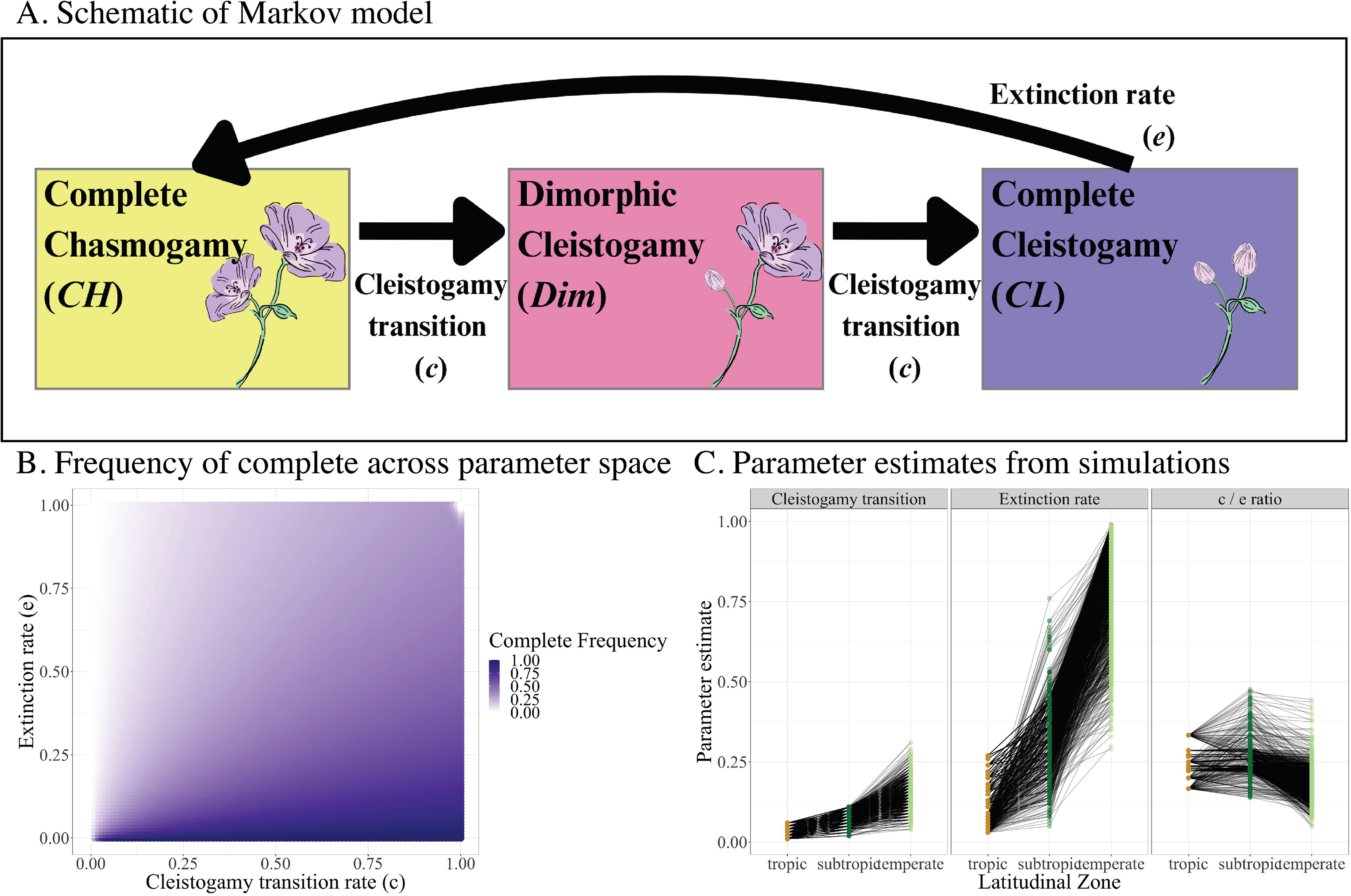
A. Schematic of Markov transition model which estimates the proportion of species that will adopt each mating strategy. There are three states: complete chasmogamy (*CH*), dimorphic cleistogamy (*Dim*), and complete cleistogamy (*CL*). Species are assumed to transition from complete chasmogamy to dimorphic cleistogamy at rate *c*, or the cleistogamy transition rate. Species also transition from dimorphic to complete at the same rate *c*. Finally, complete cleistogamous species are assumed to go extinct due to inbreeding at rate *e*, when they will be replaced by new complete chasmogamous species. B. Across the range of *c* and *e* at a step size of 0.01, we measure the frequency of complete cleistogamy from the stationary distribution. Complete cleistogamy is highest in frequency (dark blue) when the cleistogamy transition rate is high and extinction rate is low. C. Using the observed frequency of dimorphic and complete species across latitudinal zones, we perform replicate simulations to calculate the parameters for cleistogamy transition rate, extinction rate, and *c*/*e* ratio that yield the expected proportion of complete and dimorphic cleistogamy in each latitudinal zone. Black lines connect the parameter estimates for each zone within a single simulation. We assume that moving from equator to poles, both cleistogamy transition and extinction rate must increase. We find that extinction rate increases more than the cleistogamy transition rate. In the majority of simulations, the *c*/*e* ratio remains constant between tropic and subtropic zones.

We calculate the expected proportion of species in each state across all possible values of both *c* and *e*. We present the proportion of complete cleistogamous species, in order to illustrate the effect of *c* and *e* (fig. 4B). When the cleistogamy transition rate increases, complete cleistogamy increases in frequency. On the other hand, when the extinction rate increases, complete cleistogamy decreases in frequency. In other words, as the transition rate in (*c*) increases and as the transition rate out (*e*) decreases, the frequency of complete cleistogamy increases.

Given the calculated proportion of species that adopt each strategy across the *c* and *e* parameter space in our Markov model, we then estimate the parameter values for each latitudinal zone that yield the observed frequency of both cleistogamous strategies (fig. 4C). Because our model assumes that both *c* and *e* must increase as we move from the tropics to the temperate zone (Geppert et al. [2023], Ludewig et al. (2023), Baskett et al. [2020], Ollerton [2017], Mittelbach et al. [2007], Saupe [2023], Weir and Schluter [2007]) the space of possible parameter values for each zone are constrained by the parameter value for the previous zone. As such, we performed simulations where the value of *c* and *e* where randomly chosen for each zone sequentially, constrained by the parameter values of the previous zone (see methods). Parameter values for each zone from the same simulation are connected with black lines.

We estimate that the cleistogamy transition rate in the tropics ranges from 0.01 to 0.06, while the extinction rate in the tropics ranges from 0.03 to 0.27 (fig. 4C). In the subtropic zone, the cleistogamy transition rate ranges from 0.02 to 0.11 and the extinction rate ranges from 0.07 to 0.78. Despite the fact that when moving from tropics to subtropics, the range of possible extinction rates increases more than the range of possible cleistogamy transition rates, we see that in the majority of simulations the *c*/*e* ratio remains relatively constant. This likely drives the the proportion of cleistogamous species that engage in the complete strategy to remain constant in the tropic and subtropic zones (fig. 3B). In the temperate zone, the range of possible cleistogamy transition rates is from 0.03 to 0.35, while the range of possible extinction rates is from 0.32 to 0.99. Thus, in all simulations, the extinction rate increases far more than the cleistogamy transition rate when moving from subtropic to temperate zones (fig. 4C). These results imply that: a) extinction rate increases more quickly than cleistogamy transition rate, and b) the increase in extinction rate is nonlinear. While the evolution and maintenance of cleistogamy is complicated by other selective pressures, our model provides a heuristic results that differences in the relative increases in both selective pressure to self and extinction rate can recover the realized latitudinal distribution in complete and dimorphic cleistogamous strategies.

## 4 Discussion

Cleistogamy is a widespread mating system where plants produce some proportion of closed, exclusively selfing flowers. Previous research has suggested that cleistogamy may be a bet hedging strategy by reducing risk in the face of inconsistent pollinator availability and other environmental stressors (Schoen and LLoyd (1984), Berg and Redbo-torstensson [1998], Culley and Klooster [2007]). We perform a biogeographic analysis to determine where cleistogamy has evolved and been maintained, in order to evaluate the bet hedging hypothesis. We find that cleistogamy has been observed in the majority of ecoregions on earth, as well as appearing on every continent (including Antarctica, which is home to the perennial grass *Deschampsia antarctica*). We also show that cleistogamy is more commonly found in climatically variable ecoregions, and more likely to be absent in constantly warm (tropical) or cold (polar) environments. Cleistogamy is especially prevalent in Mediterranean ecoregions and temperate forests, which are both defined by high within-year seasonal variation as well as high between-year climate variation (Rundel et al. [2016], Dreiss and Volin [2020], Loidi et al. (2022)). Cleistogamy presence is also associated with lower mean temperatures, wider temperature ranges, and larger precipitation seasonality.

We also find differences in distribution between types of cleistogamous strategies. First, we investigate differences in the distribution of members of the Poaceae family and other species, which we use as a proxy for pollinator mode (Campbell et al. [1983]). Due to limitations in the scope of our dataset, we use taxon as a proxy for pollination mode. However it is likely that species outside of the Poaceae family may also be abiotic pollinated. Species that are not members of Poaceae, i.e. are primarily biotically pollinated, are more prevalent in temperate latitudes and cooler mean temperatures, consistent with the hypothesis that cleistogamy evolved in these species to hedge against variability in biotic pollinators (Ollerton [2017], Baskett et al. [2020], Oakley et al. [2007]). On the other hand, members of the Poaceae family, or abiotic pollinated grasses, are more prevalent in ecoregions with lower mean precipitation and higher precipitation seasonality, suggesting that cleistogamy in these species may insulate against variability in precipitation. Additional work elucidating the diversity in pollinator mode at the species level, and the relationship between pollinator and climate, is needed to further explain the distribution of cleistogamy.

Finally, we contrast differences between complete and dimorphic cleistogamy strategies. We find that complete cleistogamy is least prevalent in temperate latitudes, and is associated with warmer mean temperatures, narrower temperature ranges, and lower mean precipitation. We hypothesize that this is driven by the opposing pressures of selection to increase cleistogamy and extinction. Decreasing temperatures in temperate latitudes drive pollinator abundance to decrease (Geppert et al. [2023], Ludewig et al. (2023), Baskett et al. [2020], Ollerton [2017]), which likely increases selective pressure to engage cleistogamy. However, decreasing temperatures are also correlated with an increase in extinction rate (Mittelbach et al. [2007], Saupe [2023], Weir and Schluter [2007]). As complete cleistogamy leads to inbreeding, it seems likely that the increased extinction rate will cause larger decreases in the prevalence of complete cleistogamy compared to dimorphic cleistogamy. We thus hypothesize that the increase in selective pressure to self drives the frequency of dimorphic cleistogamy to increase, while the increased extinction rate causes the frequency of complete cleistogamy to decrease. We develop a a simple Markov transition model to demonstrate that increases in both the selective pressure to self and extinction rate can predict the relative abundance of complete and dimorphic cleistogamy across latitudinal zones.

So is cleistogamy a bet hedging strategy? The biogeographic distribution, specifically of dimorphic cleistogamy, shows that cleistogamy is most prevalent in environments we would expect bet hedging to evolve in. However, proving that a bet hedging trait is adaptive is more challenging. One must be able to demonstrate that the putative bet hedging trait lowers mean fitness and fitness variance within the environmental context it evolved (Simons [2011]). Classically, this has been done by assaying the geometric mean fitness of the trait (Simons [2011], Starrfelt and Kokko [2012]). However, we lack the comprehensive life history trait measurements across environments needed to calculate geometric mean fitness for the majority of cleistogamous species. Future work that measures the relative fitness of flower morphs across environmental conditions in nature is needed to answer this question.

Even in cases where fitness estimates for both flower morphs have been found, there are still issues proving that cleistogamy is adaptive. Given the low relative fitness of the chasmogamous flower morphs, empirical studies of dimorphic cleistogamy have been unable to show that the strategy increases geometric mean fitness relative to complete cleistogamy (Oakley et al. [2007]). Interestingly, recent theoretical work has found that bet hedging strategies can be more adaptive than geometric mean fitness predicts (Weissman et al. [2024], Zhou and Xue [2022]). In Weissman et al. [2024], we demonstrated that bet hedging can be beneficial at sufficiently large population sizes, even when the geometric mean fitness of the bet hedger is less than that of a resident wild-type. This suggests that dimorphic cleistogamy may be adaptive, even if it causes a decrease in geometric mean fitness relative to complete cleistogamy. As empiricists continue to assay the fitness of complete and dimorphic cleistogamy strategies across environments, it will be important to evaluate their adaptive significance using stochastic models in addition to geometric mean fitness.

Differences in the distribution of complete and dimorphic cleistogamy can help bet hedging researchers explore differences in the evolution of conservative and diversified strategies. While both conservative and diversified bet hedging adaptations have been observed in closely related taxa across the tree of life (Simons [2011]), little is known about what shapes the choice of bet hedging strategy. Attempts to model the evolution of bet hedging strategy have found conflicting results, although they often invoke different types of environmental change as a potential driver (Starrfelt and Kokko [2012], Botero et al. [2015], Einum and Fleming [2004]). However, this is complicated by the question: is complete cleistogamy actually conservative bet hedging? In some ways, the cleistogamous flower is analagous to our expectations of a conservative, low-risk phenotype: the flower exhibits no variance in fitness as pollinator availability changes. In other ways, cleistogamous flowers could be considered the specialist, high-risk phenotype, given their on average higher fitness and risk of extinction due to inbreeding. In our case, we find that complete cleistogamy is more prevalent in environments that are better on average and less variable, which is more consistent with expectation for a specialist, high-risk phenotype. Further work quantifying the relative fitness of both flower morphs across environments, with special attention to complete cleistogamous species, could help elucidate differences in the evolution of bet hedging strategy.

Another limitation of this study is the lack of climate data on between-year variability. Our finding that cleistogamy is more prevalent in ecoregions with high within-year variability (i.e. temperature range and precipitation seasonality) is consistent with previous research demonstrating that the majority of cleistogamous species exhibit seasonal germination phenology between flower morphs. However, the benefit of bet hedging is most frequently associated with lowering between-year fitness variance (Simons [2011], Starrfelt and Kokko [2012]). While we note that within-year variability in temperature is positively correlated with between-year variability in temperature, we are limited to locations within the United States where both types of climate data are available (supplemental fig. 1B). Expanding climate data to include between-year variability, as well as frequency of extreme events such as droughts, wildfires, and hard freezes, is necessary to further explore if cleistogamy is bet hedging.

The relationship between cleistogamy, other methods of mixed mating, and outcrossing rate is also an area for future work. Cleistogamy is only one method plants can engage in mixed mating; monomorphic mixed mating, or the production of a single flower morph that is capable of both selfing and outcrossing, is actually more common than dimorphic cleistogamy (Moeller et al. (2017), Goodwillie et al. [2005], Oakley et al. [2007]). Interestingly, a similar biogeographic analysis of mixed mating species revealed that mixed mating is also more prevalent in temperate latitudes (Moeller et al. (2017)). Given the energetic economy of the cleistogamous flower morphs, it has been hypothesized that dimorphic cleistogamy may be more adaptive than monorphic mixed mating when selection to maintain outcrossing is stronger (Oakley et al. [2007]). However, many dimorphic cleistogamous species have very low outcrossing rates, as even their chasmogamous species primarily self. For example, *Amphicarpaea bracteata*, an annual legume that engages in dimorphic cleistogamy, rarely reproduces via outcrossing (Kartzinel et al. [2016]). Further analysis quantifying differences in realized outcrossing rates amongst cleistogamous species, and contrasting cleistogamy with other methods of mixed mating, would provide additional context for the evolution of this strategy.

Cleistogamy is an important adaptation in angiosperms to reduce risk and cope with environmental uncertainty. This mixed mating strategy is very widespread, having evolved independently over 40 times across over a third of all angiosperm orders (Culley and Klooster [2007]). Given the relationship between cleistogamy and changing environments, gaining a deeper understanding in large-scale trends in cleistogamy can help us better understand its evolution and maintenance. Our results demonstrate a large-scale association between cleistogamy presence and variable environments, consistent with bet hedging predictions. As anthropogenic climate change causes climates to become warmer on average, more variable, and increases the frequency of extreme climate event (Change (2007)), understanding when cleistogamy, as well as similar risk reduction strategies, are beneficial can help predict species resilience.

## Supporting information

Supplemental Results

## Acknowledgements

Thank you to David Peede and Neal Yin, for your discussions on amphicarpy in *Amphicarpaea bracteata*, which have helped develop our interest and understanding of bet hedging in plants.

---

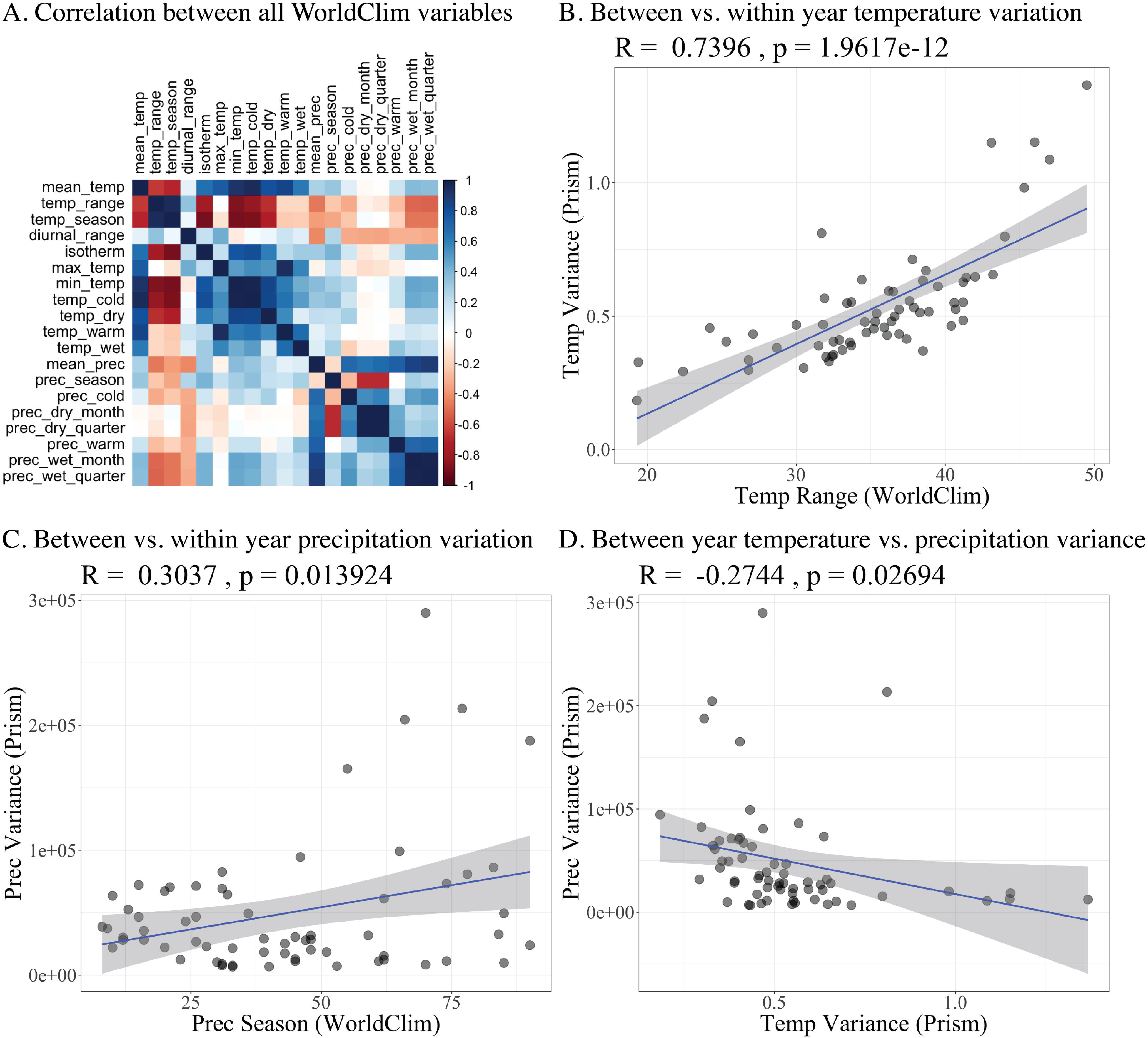

