## Supplemental Results for "Balancing the risks of mating: biogeographic evidence of cleistogamy as a bet hedging strategy"

### 1 Supplemental Results

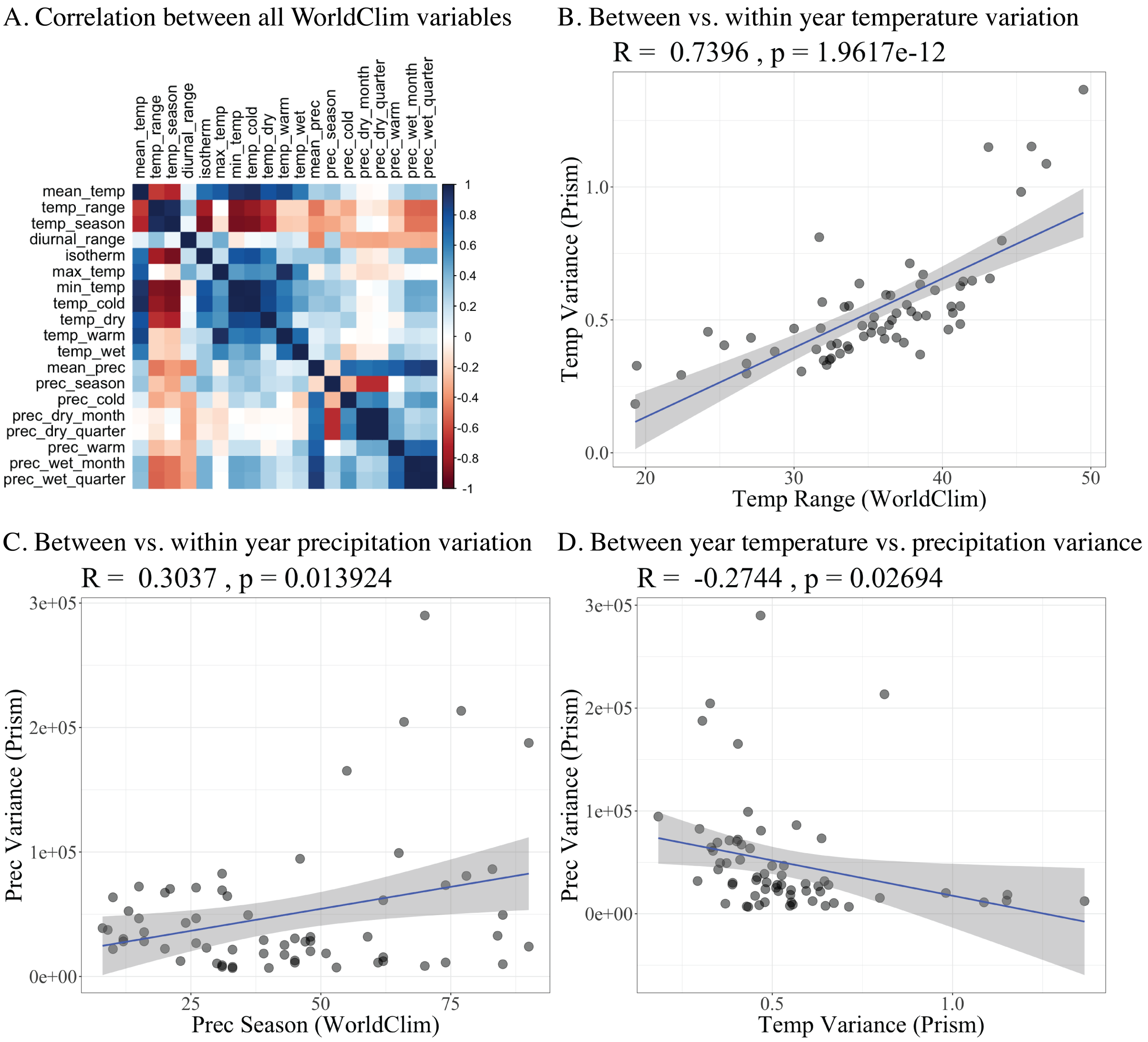


Supplemental Figure 1: A. Correlations between the 19 bioclimatic variables from WorldClim. Of the four climate variables we focus on, mean temperature and temperature range are the most highly correlated ($-0.65$). B-D. Comparing within-year variability data from WorldClim to between-year variability, calculated using 40 year annual averages, from PRISM. We use only the 65 ecoregions within the USA, as PRISM only supplies climate data from the USA. Temperature is measured in degrees Celsius, precipitation is measured in millimeters per month. B. Within-year temperature range is strongly positively correlated with between-year temperature variance. C. Within-year precipitation seasonality is weakly positively correlated with between-year precipitation variance. D. Between-year temperature variance is weakly correlated with between-year precipitation variance.

| variable | tropic | subtropic | temperate | polar |
|---|---|---|---|---|
| mean_temp | 0.0331* | 0.9336 | 0.1173 | 0.2558 |
| temp_range | 2e-04*** | 0.5551 | 0.9592 | 0.9336 |
| temp_season | 0.2558 | 0.6112 | 0.8682 | 0.8305 |
| diurnal_range | 0**** | 0.223 | 0.5252 | 0.6946 |
| isotherm | 0.7629 | 0.8682 | 0.2736 | 0.2819 |
| max_temp | 0.95 | 0.7629 | 0.0738 | 0.0036** |
| min_temp | 2e-04*** | 0.8829 | 0.2367 | 0.5055 |
| temp_cold | 0.0189* | 0.8682 | 0.2198 | 0.4597 |
| temp_dry | 0.0482* | 0.9557 | 0.8163 | 0.3371 |
| temp_warm | 0.1323 | 0.95 | 0.1013 | 0.007** |
| temp_wet | 0.0324* | 0.95 | 0.0324* | 0.2884 |
| mean_prec | 0.323 | 0.1778 | 0.989 | 0.3371 |
| prec_season | 0.0327* | 0.4813 | 0.223 | 0.5147 |
| prec_cold | 0.989 | 0.95 | 0.95 | 0.361 |
| prec_dry_month | 0.0036** | 0.7213 | 0.8946 | 0.2884 |
| prec_dry_quarter | 0.0051** | 0.6946 | 0.9336 | 0.2884 |
| prec_warm | 0.2198 | 0.2198 | 0.9336 | 0.2155 |
| prec_wet_month | 0.5789 | 0.2198 | 0.9336 | 0.3371 |
| prec_wet_quarter | 0.8682 | 0.2116 | 0.9336 | 0.323 |

Supplemental Table 1: Adjusted p-values for t-test of the difference between ecoregions where cleistogamy is present vs. absent for all WorldClim bioclimatic variables, subdivided by latitudinal zone. Significance is denoted with asterisks.

| variable | tropic | subtropic | temperate |
|---|---|---|---|
| mean_temp | 0.0914 | 0.1582 | 0**** |
| temp_range | 0.2758 | 0.2758 | 0.0089** |
| temp_season | 0.5579 | 0.4906 | 0**** |
| diurnal_range | 0.3011 | 0**** | 1e-04**** |
| isotherm | 0.4906 | 0.0017** | 0**** |
| max_temp | 0.0314* | 0.0132* | 0.0285* |
| min_temp | 0.3679 | 0.475 | 0**** |
| temp_cold | 0.1387 | 0.0914 | 0**** |
| temp_dry | 0.1464 | 0.0882 | 0**** |
| temp_warm | 0.0918 | 0.2427 | 0.3011 |
| temp_wet | 0.138 | 0.0067** | 0.0024** |
| mean_prec | 0.4572 | 0**** | 0.941 |
| prec_season | 0.3211 | 0.3939 | 0.4548 |
| prec_cold | 0.9018 | 1e-04**** | 0.0177* |
| prec_dry_month | 0.3121 | 0.0016** | 0.2397 |
| prec_dry_quarter | 0.3298 | 0.0016** | 0.3679 |
| prec_warm | 0.2397 | 0**** | 0**** |
| prec_wet_month | 0.4906 | 0**** | 0.9018 |
| prec_wet_quarter | 0.5668 | 0**** | 0.9018 |

Supplemental Table 2: Adjusted p-values for t-test of the difference between occurrences of complete vs. dimorphic cleistogamy for all WorldClim bioclimatic variables, subdivided by latitudinal zone. As there are no occurrences of complete cleistogamy within the polar zone, the polar zone is not included. Significance is denoted with asterisks.

| var | complete | dimorphic |
|---|---|---|
| mean_temp | 7e-04*** | 0**** |
| temp_range | 0.1466 | 0.0524 |
| diurnal_range | 1e-04**** | 0**** |
| isotherm | 0.2417 | 0**** |
| max_temp | 0**** | 0**** |
| min_temp | 0.0617 | 0**** |
| temp_cold | 0.0105* | 0**** |
| temp_dry | 0.0023** | 0**** |
| temp_season | 0.5813 | 0**** |
| temp_warm | 0**** | 0.0034** |
| temp_wet | 5e-04*** | 0.774 |
| mean_prec | 0**** | 0**** |
| prec_season | 0**** | 0.0691 |
| prec_cold | 0**** | 0.0018** |
| prec_dry_month | 0**** | 0**** |
| prec_dry_quarter | 0**** | 0**** |
| prec_warm | 1e-04**** | 0**** |
| prec_wet_month | 0.0023** | 0**** |
| prec_wet_quarter | 5e-04*** | 0**** |

Supplemental Table 3: Adjusted p-values for t-test of the difference between occurrences of members of the Poaceae Family (wind pollinated) vs. members of all other Families (primarily biotic pollinated) for all WorldClim bioclimatic variables, subdivided by cleistogamy type. As both pollination types have similar latitudinal distributions, we do not subdivide data by latitudinal zone. Significance is denoted with asterisks.
